# Phylogenomics of Asgard archaea reveals a unique blend of prokaryotic-like horizontal transfer and eukaryotic-like gene duplication

**DOI:** 10.1101/2025.07.24.666526

**Authors:** Saioa Manzano-Morales, Toni Gabaldón

**Affiliations:** Barcelona Supercomputing Center (BSC). Plaça Eusebi Güell, 1-3 08034 Barcelona, Spain; Institute for Research in Biomedicine (IRB Barcelona), The Barcelona Institute of Science and Technology, Baldiri Reixac, 10, 08028 Barcelona, Spain; Catalan Institution for Research and Advanced Studies (ICREA), Barcelona, Spain; CIBER de Enfermedades Infecciosas, Instituto de Salud Carlos III, Madrid, Spain

## Abstract

Asgard archaea hold a pivotal position in the tree of life as the closest known relatives to eukaryotes and are therefore crucial for understanding eukaryogenesis. Earlier genomic analyses revealed that Asgard genomes are remarkably larger than those of other archaea and contain a significant number of genes seemingly acquired from bacteria. However, the precise contributions of horizontal gene transfer and gene duplication in shaping Asgard genomes remain largely unknown. Here, we present a comprehensive phylogenomic analysis to dissect the evolutionary dynamics of Asgard genomes, quantifying gene duplication, loss, and both inter- and intra-domain gene transfer events. Our findings reveal that gene transfer is widespread throughout Asgard evolution, predominantly affecting metabolic genes at the periphery of interaction networks. However, our analyses demonstrate that gene duplications, rather than horizontal gene transfers, are the primary drivers behind the increased genome sizes observed in Asgard archaea. This unique evolutionary signature in Asgard archaea—a blend of pervasive prokaryotic-like gene transfer alongside significant eukaryotic-like gene duplication—is consistent with their phylogenetic placement and offers novel insights into the genomic transitions that likely underpinned eukaryogenesis.

## Introduction

Asgard archaea are the closest prokaryotic relatives of eukaryotes and are therefore key to understanding eukaryogenesis^1,2^. Asgard genomes encode many of what were previously thought to be eukaryote-specific proteins^2^, including actin and actin-related proteins^3^, and an actin cytoskeleton has been found in a cultivated Lokiarchaeum^4^. The crucial interest of Asgard archaea has promoted sequencing efforts, resulting in a large number of available genomes - 296 assemblies in release 226 of the Genome Taxonomy Database (GTDB)^5^. These represent 12 class-level lineages *sensu* GTDB (namely Asgard-, Atabey-, Baldr-, Heimdall-, Hermod-, Jord-, Loki-, Njord-, Odin-, Sif-, Thor-, and Wukongarchaeia). However, most data corresponds to Metagenomic Assembled Genomes (MAGs), which can be contentious due to potential incompleteness or contamination issues^6^ . Asgard archaea have proven elusive to culture, with two exceptions from Lokiarchaea: *Ca.* Prometheoarchaeum syntrophicum MK-D1^7^ and *Ca.* Lokiarchaeum ossiferum B-35 4, which can be cultured with their syntrophic partners and therefore have enabled high-quality genomes.

Asgard archaea were first identified in a deep-sea hydrothermal vent^8^, but their presence in clone libraries precedes their description by two decades^9–11^. Recent sampling efforts have uncovered new Asgard lineages, of which most were first described in anoxic marine environments^12^. However, metagenomic analyses indicate a worldwide cosmopolitan distribution^13^. This reflects on a broad array of lifestyles and metabolic properties including mixotrophy, homoacetogenesis, facultative anaerobiosis, alkane utilization, and rhodopsin-based phototrophy^13–17^. However, we have a very limited understanding of how genome variation underlies such diverse adaptations.

Earlier analyses of Asgard genomes uncovered a large number of putatively transferred bacterial genes^18^ and generally larger genome sizes as compared to other archaea^19^. Previous analyses considering prokaryotes have found weak^20^ to strong^21,22^ positive correlations between genome size and horizontal gene transfer (HGT) fraction, but none of them included Asgard genomes. Similarly, Asgard archaeal genomes have been shown to have higher rates of duplication than other archaea^19^, but the functional implications of these gene duplications, or how these duplication tendencies contrast with HGT (the prevailing form of genome expansion in prokaryotes) have not been comprehensively explored .

Here, we reconstructed the Asgard archaeal pangenome and used phylogenomics on a super-phylum-wide scale to assess the relative contribution of gene duplication, loss and transfer in the evolution of Asgard genomes against the backdrop of its sister clade, Thermoproteota, and investigate possible functional trends. With this we aimed to shed light on the processes that have specifically driven Asgard archaea towards larger genome sizes and other ‘eukaryote-like’ qualities in contrast to Thermoproteota, which remain more ‘prokaryote-like’. Our results indicate that Asgard archaea have high duplication rates that are intermediate between those in their prokaryotic sister clade (despite similar transfer rates) and those observed in unicellular eukaryotes. We uncover widespread but mostly lineage-specific inter-domain HGT involving several donor-acceptor pairs and metabolic genes. Overall, our results indicate that Asgard genomes are uniquely shaped by a blend of prokaryotic horizontal gene transfer and eukaryotic-style gene duplications.

## Results

### The pangenome and signature proteins of Asgard archaea

We first sought to characterize the Asgard pangenome by performing homology-based clustering of all available genomes from phylum Asgardarchaeota in GTDB^5^. However, nearly all Asgard genomes are MAGs, resulting in a median CheckM^23^ completeness of 87.54% (SD 8.21). This lack of genome completeness can lead to an infra-estimation of the core genome and, conversely, an over-estimation of the accessory genome.

For this reason, we compiled a representative subset of 32 genomes covering all family-level lineages of Asgard archaea by choosing the highest-quality available genome (following the GTDB score criterion) per family and prioritizing available isolates and closed genomes (See Methods, Supplementary Data 1). Additionally, we included 163 other prokaryotic genomes as out-groups, including 28 representative species from the Asgard archaea’s sister group, TACK (“Thermoproteota” hereafter, *sensu* GTDB). We clustered proteins into families (orthogroups, OGs) using a similarity-based approach (See Methods). This yielded a total of 71,402 OGs, including singletons, out of which 17,429 contained at least one Asgard protein. Of these, 12,397 were exclusively present in Asgard archaea, and of these 10,470 were present in a single Asgard genome and no other prokaryote in our dataset (singletons). 79 gene families were found in all Asgard genomes (strict core) and 310 in 90% or more (relaxed core, see Figure 1A for the distribution), with the remaining belonging to the accessory pangenome. We annotated OGs with KEGG Orthologs (KOs, see Methods) and found that proteins in the Asgard relaxed core were enriched in pathways such as ribosome (ko03010), DNA replication (ko03030), and aminoacyl-tRNA biosynthesis (ko00970) (Supplementary Data 2).

**Figure 1:**
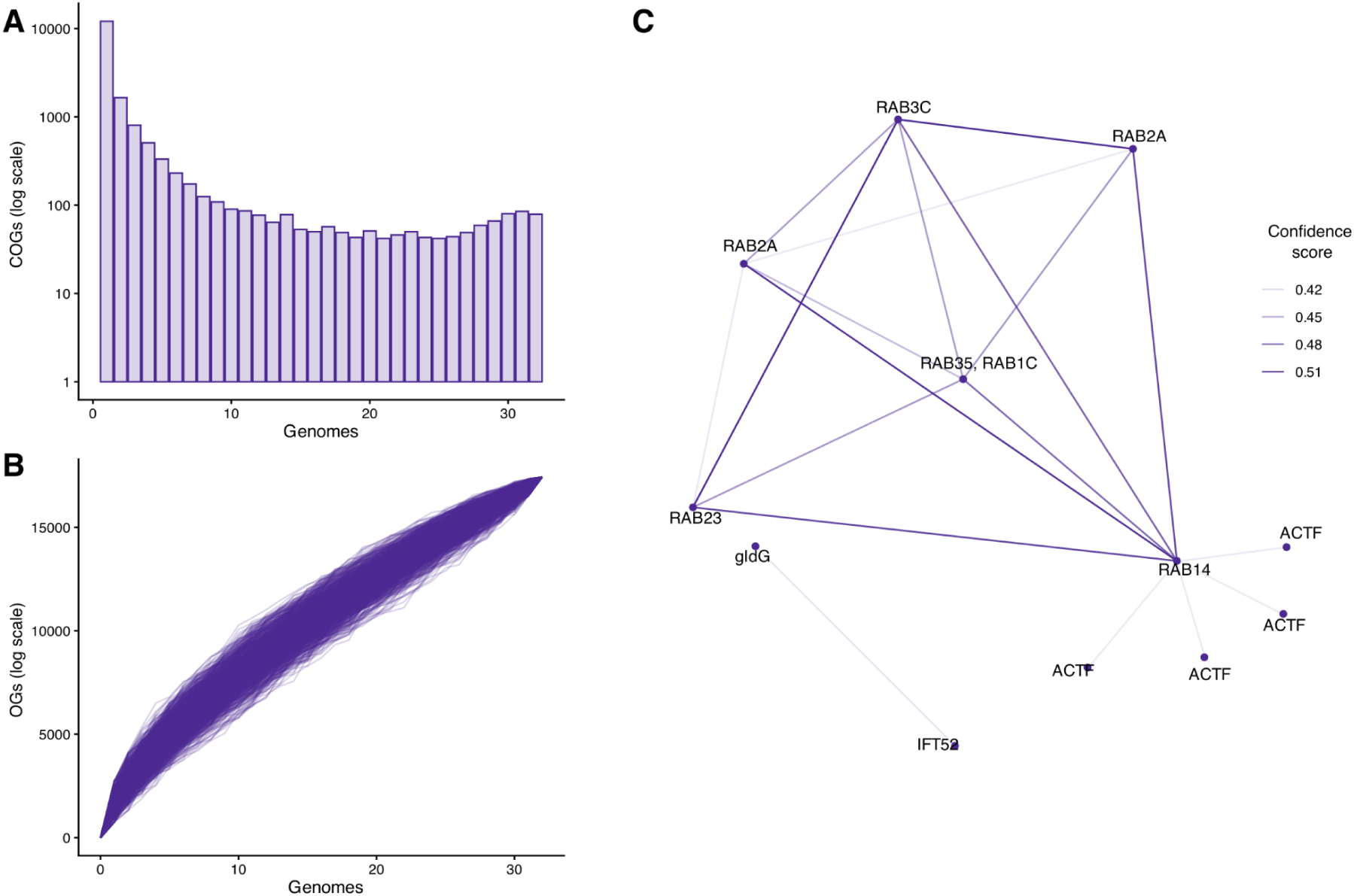
Asgard archaeal pangenome metrics for the high-quality subset. (A) Frequency of OGs according to the number of genomes in which they are present in the pangenome of Asgard archaea (bin width=1). (B) Rarefaction curves obtained by subsampling the genomes in a random order over 1000 iterations. (C) Protein-protein interaction network of the Asgard-Signature proteins. Each node is one non-redundant protein (labeled as its annotated KO/KOs), edges represent protein-protein interactions with color strength proportional to the interaction score given by STRING.

We estimated the size of the total Asgard pangenome both by Chao’s lower bound^24^ (yielding a lower bound estimate of 61,815 OGs) and by the best-fitting binomial mixture model (resulting in a pangenome of 75,659 OGs and a strict core of 18 OGs). We additionally calculated rarefaction curves (Figure 1B) which, along with the Heaps estimate (alpha = 0.26), indicated an open pangenome.

To assess the impact of MAG incompleteness on pan-genome size estimates, we repeated the analyses replacing the 32 best-quality representatives by 32 randomly-chosen, taxonomically-equivalent genomes. In this case, we obtained 72,179 OGs, with 18,135 including at least one Asgard representative, and a strict and relaxed observed core of 57 and 269, respectively. We inferred a similarly open pangenome estimate with identical Heaps alpha (0.26) and strict core inferred by the binomial mixture model (18), but a higher Chao’s lower bound (72,712) and total estimated size by the binomial mixture model (94,783). Hence, both genomic subsets provided similar estimates in terms of openness, OGs and total pan-genome size but not in terms of observed core genome or lower bound estimates, which were more affected by genome incompleteness.

As the core size estimate is particularly sensitive to genome incompleteness^25^, we ran mOTUpan^26^ for a more robust estimation of core size and fluidity, as this approach intrinsically considers genome incompleteness (as provided by CheckM completeness). This approach estimated a larger core set for both of the high-quality (473) and random (509) datasets. Hence, even when considering the most conservative estimates, these results underscore how incomplete is our knowledge of the breadth of Asgard archaeal protein diversity, implying the intriguing possibility that new protein families with potential implications for eukaryogenesis may await discovery.

We defined Asgard signature proteins (ASPs) as those OGs common (present in ≥ 86% of the proteomes, see Methods for the choice of threshold) in Asgard archaea but rare (present in ≤15% of the proteomes, following the ‘cloud’ pangenome definition, see ref.^27^) in other archaea or bacteria. This rendered 14 ASPs, for which we obtained a consensus of the annotations across Asgard proteomes and found them to be enriched in processes such as endocytosis (KEGG pathway ko04144) or motor proteins (ko04814), see Supplementary Data 2). We cross-mapped a set of 83 InterPro domains associated with 79 Eukaryotic Signature proteins (ESPs) recently identified in Asgard archaea^19^ and found 44 out of 79 of them in our dataset. We found all ASPs within the set of OGs comprising ESPs, suggesting that these eukaryotic-like proteins contributed to the differentiation of Asgard archaea from other prokaryotes and that some of these differentiating gene families ended up into the eukaryotic lineage. ASPs include proteins previously considered ESPs involved in typical eukaryotic processes such as Ras-related proteins (K07877), ESCRT-II complex subunit VPS22 (K12188), actin (K10355) and actin-related proteins (K10369), as well as other motor proteins (K10398). In addition to these ESPs, ASPs include other relevant proteins such as motility proteins (K25153) which are putative adaptations to a mat environment^28^.

All 14 ASP OGs are found in *Ca.* Prometheoarchaeum syntrophicum MK-D1, where they comprise 90 proteins. We used this organism’s protein-protein interaction network (PPI) in STRING^29^ as a template to assess the connectivity of these ASPs (Figure 1C), which indicated that they are more connected than would be expected by chance (PPI enrichment p-value of 5.32e-09).

To identify lineage-level signature proteins that may underlie lineage-specific adaptations, we selected OG that are part of the lineage’s core and rare outside (cloud-level, ≤15%). These lineage-signature families were enriched in functions such as “two-component system” (ko02020) in Sifarchaeia, and “ABC transporters” and “Various types of N-glycan biosynthesis” (ko02010, ko00513) in Lokiarchaeia (Supplementary Data 2).

### Genome dynamics across Asgard archaea

To assess dynamic changes and their underlying mechanisms in Asgard genomes, we performed a comprehensive phylogenomic analysis. For this, we ran a species overlap^30^ and tree reconciliation^31^ pipelines (see Methods).

For the species overlap pipeline, we selected ten Asgard and ten Thermoproteota representatives and used them as seeds for the reconstruction of the evolutionary histories of all their proteins in the context of the 60 Asgard+Thermoproteota genomes, such that each phylum was balanced in the dataset. These 20 phylomes comprised a total of 38,889 gene phylogenies (available in PhylomeDB^30^ as phylomes 1532-1551).

For the reconciliation pipeline, we reconstructed the gene trees of the amino acid sequences of the orthogroups from OrthoFinder that had sequences from at least four species, and included at least one Asgard/Thermoproteota sequence, totaling 4,699 families and 344,709 proteins.

We then performed reconciliation and species-overlap analyses using two alternative species tree topologies for the Asgard clade due to ongoing debates (see Methods) to i) detect and date gene duplication events, including gene family expansions ii) infer orthology and paralogy relationships, and iii) infer HGT. Although the reconciliation analysis was performed for the full, 195-genomes dataset, we only show results for the clades of interest, Asgard archaea and Thermoproteota.

The average number of duplications per gene and lineage (duplication rate hereafter, measured as the total number of duplications at a given node divided by the number of gene families present at that node), as computed with either the species-overlap or reconciliation algorithms were significantly correlated (p-value 4E-14, R=0.82), and we show reconciliation-based duplication rates, together with transfer and gene loss rates in Figure 2A for the Eme et al. topology (results for the alternative topology are shown in Supplementary Figure 1, which are by and large congruent).

**Figure 2:**
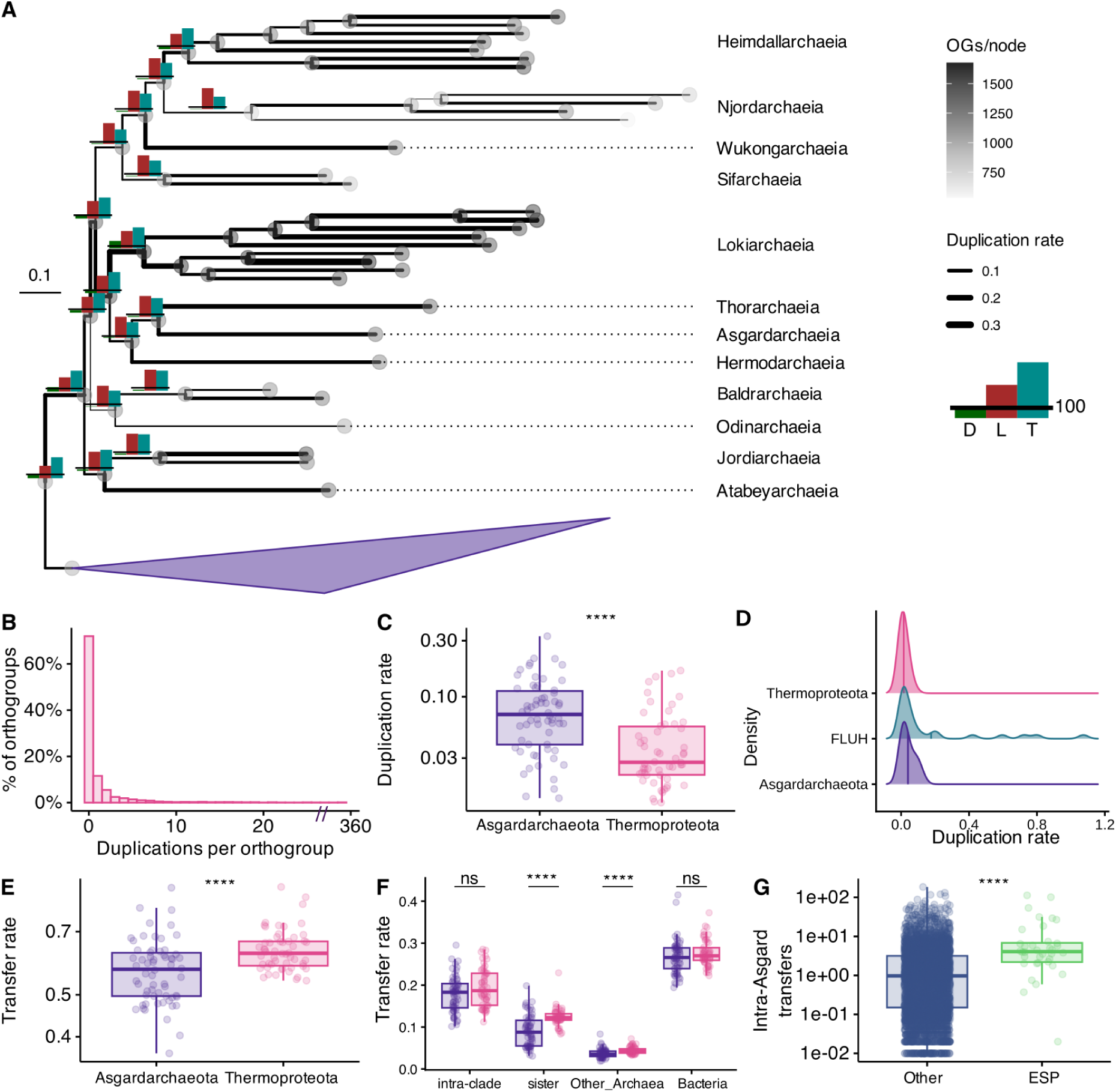
Gene gain-loss dynamics across Asgard archaea. (A) Gain and loss dynamics in Asgard archaea. The total number of OGs per proteome is displayed by circle color in each node. Duplication rate (number of duplication events relative to the inferred number of families present in the node) is represented by the branch width. Number of duplications (green), transfers (blue) and losses (red) displayed at internal nodes in absolute number. Horizontal bar indicates 100 events as scale. (B) Histogram of sum of duplication events per orthogroup across Asgard nodes. Axis cut at 30, we artificially display the tick as the highest value rounded to the multiple of 10. (C) Duplication rates per node of Asgard (n=62) and Thermoproteota (n=54) gene families, statistical significance assessed with two-tailed Wilcoxon rank test, p-value 3.3762E-06. (D) Duplication rates, inferred by species overlap, of Asgard, Thermoproteota and FLUHs. (E) Transfer rates per node of Asgard (n=62) and Thermoproteota (n=54) gene families, statistical significance assessed by two-tailed Wilcoxon rank-sum test, p-value 1.13123E-04 (F) Transfer rates per node in Thermoproteota (n=54) and Asgard archaea (n=62), separated by donor, statistical significance assessed by pairwise two-tailed Wilcoxon rank-sum tests, p-values 0.107 (intra-clade), 3.92E-07 (sister), 1.28E-05 (other archaea), 0.14 (bacteria) (G) Intra-Asgard transfer events on ESPs compared to the rest of gene families (n=36, 4295 gene families, respectively). Permutation test, p-value 2.2E-16. In the plots with significance values, asterisks * indicate p-values, * (cut-off 0.05) ** (0.01), *** (0.001) **** (0). For the box plots in 2C, E-G, in each boxplot, the central line indicates the median, the lower and upper limits of the rectangle indicate the first and third quartiles, respectively and the whiskers extend to the minimum and maximum value of 1.5 times the interquartile range.

Our results for both topologies confirmed previously described high levels of duplications at the base of Heimdallarchaeia and Lokiarchaeia^19^, but extended this observation throughout the Asgard phylogeny. Focusing on the main topology, which we will discuss hereafter except explicitly mentioned, duplication rates were particularly high at the base of the Asgard group and at the branch separating the majority of Asgard lineages from the earliest-diverging clades (that is, excluding Baldr-, Odin-, Jordi- and Atabeyarchaeia), with additional hotspots at the base of the above-mentioned Lokiarchaeia, as well as Heimdallarchaeia, a key lineage to eukaryogenesis as current reconstructions suggest eukaryotes either stemmed from within^19^ or as sister to^32^ this clade. Some of the bursts of duplications coincide with the origin of clades with larger genomes, such as Lokiarchaeia and Heimdallarchaeia, hinting towards gene duplication driving increased genome sizes in Asgard archaea. Conversely, lineages with smaller genome sizes are associated with high rates of gene loss. In many cases, gene transfers add to the increase in the genome size, uncompensated by gene loss, such as the base of Asgard, and in many other early nodes. The exceptions to this are at the Heimdall-Njord-Wukong-Sif clade, where most nodes (to the exception of the MRCA of the sampled Heimdallarchaeia) are dominated by gene loss. This result was consistent with the alternate topology.

Importantly, the high duplication rates in Asgards were the result of large expansions in a few gene families (for the OGs for which there is duplication information, 23.87% of the genes have duplicated at least once within the Asgard lineage, see Figure 2B and Supplementary Figure 1B, where the result is overall similar), which were functionally diverse. Assessing by COG category, the categories with more than 10 OGs and the highest mean duplications across Asgard were Intracellular trafficking, secretion, and vesicular transport (U, mean 11.12, sd 59.87), mobilome (X, mean 6.68, sd 18.90), and transcription (K, mean 4.08, SD 12.68). The high values of duplications for OGs associated with the mobilome indicate the genome size increase associated with gene duplication may be driven, at least partially, by these mobile genetic elements. Despite high numbers of transfers throughout the tree, and despite high variability, compared with Thermoproteota, Asgards displayed significantly higher levels of gene duplication (as mentioned in^30^, Figure 2C also true for Supplementary Figure 1C), suggesting that they might display duplication levels more akin to those in eukaryotes. To test this, we calculated the gene phylogenies of all the genes in a set of 4 eukaryotic seed genomes against a dataset of 30 Free-Living Unicellular Heterotrophic eukaryotes (FLUHs) and calculated duplication rates for FLUHs, Asgards and Thermoproteota with the species-overlap algorithm (Figure 2D, Supplementary Figure 1D). Asgard archaea displayed duplication rates that were intermediate between those in Thermoproteota and free-living unicellular eukaryotes for both topologies. In this regard, it is also notable that proteins of Asgard origin are the ones that duplicated the most in the proto-eukaryotic lineage^33^.

We wondered then if the evolutionary regime governing the evolution of Asgard genome size was more ‘prokaryote-like’ (deletion-driven) or ‘eukaryote-like’ (neutral) (Supplementary Figure 2, Methods).

Compared to eukaryotes, prokaryotes have a narrower size range for their genomes, and genome size increases correlate with increases in protein number^34^, whereas in eukaryotes such increases seem to be driven by ncDNA, especially repetitive elements^35,36^. To maintain a streamlined genome, prokaryotes couple gene transfer with extensive gene loss^34^. In both Asgard and Thermoproteota (but more so in Asgard), the number of duplications is positively correlated with the number of gene family copies (Supplementary Figure 2A), suggesting a role of gene duplication in increasing genome size, whereas we detect no statistically significant correlation between gene gains (duplications and transfers) and gene losses in either group (Supplementary Figure 2B), nor any statistically significant change in coding density in their genomes (Supplementary Figure 2C). However, when analyzing the copy number of gene families that contained integrase domains (see Methods), indicative of mobile genetic elements, we saw that, on average, Asgard nodes contain a higher copy number of these elements both in terminal and ancestral nodes (Supplementary Figure 2D), which may be indicative of their accumulation and therefore of a more neutral regime. Asgard archaea are low-abundance^2^ and with doubling times in the range of weeks^7^, which suggests a low effective population size (Ne) that may weaken selection towards keeping a streamlined genome.

Results for the duplication rate analysis with the species-overlap algorithm can be seen in Supplementary Figure 3. These results corroborate the aforementioned bursts of gene duplication in the Lokiarchaeia and Heimdallarchaeia clade, as well as in the deeper nodes of the Asgard clade.

Next, for clades with more than 2 representative genomes (to have more than 3 nodes of reference), we looked at the set of “lineage signature proteins” described above to understand the relative contribution of gene duplication to lineage diversification. We compared the duplication rates of these OGs within and outside the lineage, normalized by the overall duplication rate of that node, and found that, these signature OGs were more duplicated within than outside the lineage, although it was only significant for Lokiarchaeia (Supplementary Figure 4), hinting to a possible role of gene duplication in lineage-specific adaptations in these lineages

We searched for tandem duplications (e.g. resulting in sets of paralogous genes that are contiguous in the genome), and found a highly variable number across Asgard genomes (Supplementary Figure 5A). Closer inspection of tandem duplications, revealed interesting cases of the same gene, or block of genes, being repeatedly duplicated in tandem (which in some cases were preserved across several genomes), as well as cases of different tandem duplications occurring contiguously in the genome (Supplementary Data 3). We performed a KEGG pathway enrichment for genes within tandem duplications and found them to be enriched in several metabolic functions including ABC transporters, butanoate metabolism, carbon metabolism, endocytosis, Microbial metabolism in diverse environments, and quorum sensing (Supplementary Figure 5B, Supplementary Data 4). Paralogs within tandem duplications had signs of purifying selection (median dn/ds of 0.367, sd 0.137, n = 8694), suggesting both copies remain functional.

### Horizontal Gene Transfer in Asgard archaea

We then assessed HGT, both intra- and inter-domain, across Asgard archaeal genomes. We first analyzed HGT using gene tree reconciliation algorithms in the above described phylogenetic dataset (see Methods). On average, and despite high variability, Asgard archaea have significantly lower transfer rates than its sister clade (Figure 2E, Supplementary Figure 1E). This difference is mostly driven by lower rates of acquisition of genes in Asgards from other Archaeal donors, including Thermoproteota, whereas acquisitions from Bacterial donors were similar for the main topology (Figure 2F) but slightly higher for Asgard archaea in the alternate topology (Supplementary Figure 1F). Among Asgard archaea, for lineages with more than 1 representative, all lineages had similar levels of HGT (Kruskal-Wallis chi-squared =10.082, df = 5, p-value =0.07295), while the duplication rates were lineage-dependent (Kruskal-Wallis chi-squared =19.472, df = 5, p-value = 0.001569).

We focused on intra-Asgard HGT exchange partners. HGT patterns depict a dense network, in which most partners have exchanged a comparatively low number of genes, but where certain key pairs stand out (Figure 2A and Supplementary Figure 6), such as highways between Loki and Heimdall clades. For the transfer pairs beyond percentile 95, we ran a KEGG pathway enrichment of the orthogroups transferred with probability ≥ 0.7 and obtained an enrichment in pathways related to metabolic functions (“Glycolysis / Gluconeogenesis”, “Propanoate metabolism”, “Porphyrin metabolism”, “Degradation of aromatic compounds”, “Pyruvate metabolism”, “Butanoate metabolism”, “Microbial metabolism in diverse environments”, “Benzoate degradation”, and “ABC transporters” were the pathways enriched in at least 2 pairs). Overall, we interpret these results as evidence of a high degree of phylogenetic incongruence in the context of a strong phylogenetic backbone, with frequent and dispersed HGT. Importantly, intra-Asgard HGT has been proposed as a potential factor explaining the absence of ESPs in Heimdallarchaeia, the closest Asgard relatives of Eukaryotes^37^. In this regard, we observed that, for OGs that are transferred within Asgard nodes, ESPs had a significantly higher number of transfers as compared to other genes (Figure 2G).

To assess inter-domain HGT we performed an exhaustive search restricted to fourteen seed Asgard genomes, by using a combined BLAST- and phylogeny-based approach. as well as using a broader database comprising 491,850 sequenced genomes from prokaryotes, eukaryotes and viruses (see Methods).

Blast-based approaches suggested a large amount of HGT, involving between 11.37-19.36% (SD 2.37) of the total gene repertoire (Supplementary Data 5). As many blast-based HGT predictions are likely to be false positives, we reconstructed phylogenies for each candidate and ran two different phylogeny-based HGT detection algorithms (see Methods). This approach predicted 3,000 gene transfer events in the 14 investigated genomes, involving 1.94-13.10% (SD 3.56) of the gene repertoire. These HGT events involved 980 OGs, out of which 922 were in the AleRax dataset, that, on average, had both a significantly higher number of duplications (Wilcoxon Rank Sum Test, two-tailed, W(922,3752)=2336359 p-value 1.04E-67) and transfers (permutation test, p-value 2.2E-16) within Asgard nodes, highlighting the complex history of these gene families. The overall good agreement between the two phylogeny-based HGT detection approaches, the generally high phylogenetic support (Supplementary Figure 7A), the confirmation of a previously described HGT event^38^, and the lack of correlation with genomic contamination estimates (Supplementary Figure 7B), suggested that our pipeline is rather conservative and that our phylogeny-based inter-domain HGT inferences are likely to be correct. We additionally kept a subset of transfer events for which the bootstrap was well-supported (≥70), which we labeled as the “high-confidence” set, to test that the qualitative conclusions held at higher stringency.

Based on the gene tree topologies, we inferred the directionality of the transfers (Supplementary Figure 7C), which now allows us to discern between Asgard-to-Bacteria or Bacteria-to-Asgard transfers. Although our pipeline was designed to primarily search for Bacteria-to-Asgard HGT, we captured some Asgard-to-Bacteria transfers. We used the last common ancestor of the genomes (see Methods) carrying the transferred genes as a proxy for the relative time of the event, and mapped this onto the Asgard species tree (Figure 3A).

**Figure 3:**
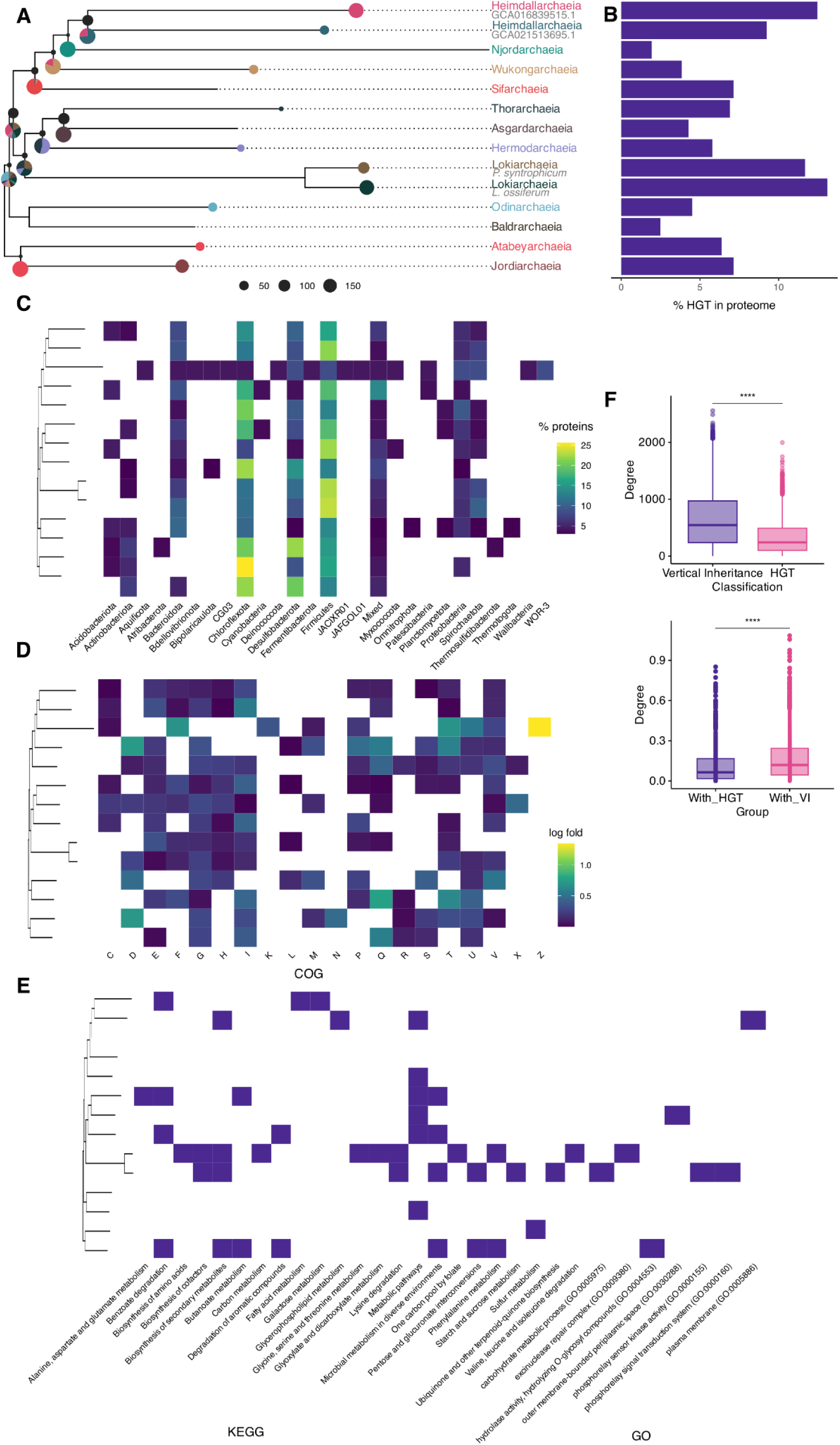
HGT-derived proteins across the Asgard lineage. A) Prevalence of HGT across the Asgard phylogeny. Dot size represents the mean number of gene trees with HGT events mapped as having occurred in that given ancestor, average of the HGT events detected per seed. Pie chart below the dots represents relative contribution of the different seeds to the inferred HGT events. Each seed (tip) proteome has a color. B) Percentage of genes involved in HGT, relative to the total number of protein-coding genes per lineage. C) Main partner phyla per lineage that have contributed at least 3% of the HGT genes to the given lineage (D-E) KEGG, COG, and GO term enrichment for the proteins derived from Bacteria-to-Asgard HGT. (F) Network properties of HGT-acquired genes versus vertically inherited ones: degree (number of interactions) of the proteins of HGT origin (n=2013) and of VI origin (n=23443, two-tailed t-test, p-value < 2.22E-16) (above); for the HGT proteins, we show the number of connections per node of the subnetwork composed only of HGT proteins (with_HGT) and calculate the number of connections with VI proteins (n=2013 nodes, two-tailed Wilcoxon rank-sum test, p-value < 2.22E-16). Degrees are normalized by the number of proteins in the protein of each category. In each boxplot, the central line indicates the median, the lower and upper limits of the rectangle indicate the first and third quartiles, respectively and the whiskers extend to the minimum and maximum value of 1.5 times the interquartile range.

We assessed the relative contribution of the different bacterial clades as exchange partners (irrespective of HGT directionality), Our results (Figure 3C) show many different partners, with varying levels of taxonomic width. Interestingly, some of the identified partners belong to taxonomic groups that are known to co-occur or interact with extant Asgards. For example, Lokiarchaeota co-occurs with Deltaproteobacteria (phylum Proteobacteria) and Anaerolineaeae (phylum Chloroflexota), both of which appear as significant transfer partners (>5% of trees). Desulfobacterota (*sensu* GTDB) is particularly interesting in the Lokiarchaeia lineages, as both *Ca.* P. syntrophicum and *Ca.* L. ossiferum have known syntrophic partnerships with such bacteria, particularly of the *Desulfovibrio* and *Halodesulfovibrio* genera^4,7^. Importantly, most of the Desulfobacterota HGT partners identified by our pipeline belong to lineages different from the co-cultured symbionts, discarding contamination as a potential source.

To understand the impact of HGT in Asgard evolution, we focused on bacteria-to-Asgard transfers in subsequent analyses. A functional assessment of the transferred genes further supports the lineage-specificity of the transfers, with some trends in the broad sense (see COG categories in Figure 3D) but with all enriched GO terms displaying a patchy distribution (Figure 3E). Similar results were obtained when using the high-confidence set, and the same applies for KEGG functional categories (Figure 3E), despite the general KEGG category “Metabolic pathways” being a common enrichment. This is in line with previous observations in prokaryotes as a whole, where ancient transfers have been found to be enriched for genes involved in amino acid, carbohydrate and energy metabolism^20^. As for COG categories, which allow a broader overview, some key categories stand out such as Z (cytoskeleton) for Njordarchaeia, T (signal transduction mechanisms) and Q (secondary metabolites biosynthesis, transport and metabolism) and I (lipid transport and metabolism) across several lineages.

ThyX, previously identified as an HGT event in Lokiarchaeia^38^, is confirmed in this analysis, as “one carbon pool by folate” is 4.31 fold enriched in *Ca.* P. syntrophicum and is also enriched in the Hodarchaeales (GCA016839515.1), Heimdallarchaeia (GCA021513695.1), Thorarchaeia (GCA_004524565.1) and Jordiarchaeia (GCA037305845.1) representatives as well as in *Ca.* L. ossiferum (GCA025839675.1), despite none of these remaining significant after FDR correction.

We next asked whether HGT-derived proteins were integrated in the recipient organism interactome. To do so, we calculated the proteome-wide protein-protein interactions using STRING^29^ and assessed the topological properties of the resulting interactome. We found that, within this network, HGT-derived proteins have a smaller degree, that is, they were less connected than proteins of vertical descent, and were less central in the network (Figure 3F). This was true for all lineages, and for other centrality measures (Supplementary Figure 8A,B).

We found that HGT-derived proteins tend to be more connected to proteins of vertical origin than to other HGT proteins, when normalizing by their respective abundance (Figure 3E). In fact, we found that the KOs brought about by the HGT events are to some extent functionally redundant, as on average, 40.93% (SD 11.66) of the HGT-derived KOs are also found in a gene that is vertically inherited in the same genome. We also estimated the metabolic coherence of these HGT proteins by estimating the metabolism that could be completed with just these HGT KOs with anvi’o (anvi-estimate-metabolism). On average, per genome, the modules that can be assembled from these HGT KOs have a mean pathwise completeness of 0.1955 (SD 0.0438). Therefore, we conclude that HGT does not generally involve the addition of whole modules, but rather a stepwise addition of enzymes to native pathways. Although this was the overall trend, in some cases HGT-derived genes formed a stepwise complete module, thereby expanding the metabolic potential of the recipient organism: it is the case for M00046 in GCA016839515.1, M00632 in *L. ossiferum* and M00086/M00045 in *P. syntrophicum*. Nevertheless, given the overlap of KOs across pathways and the uncertainty of functional inferences, claims for contributed metabolic potential should be considered with caution.

We searched for “HGT islands”, that is, segments of the genome in which more than one HGT gene is located contiguous to another HGT gene or interspaced by a single vertically inherited gene. Most of the HGT genes are interspaced with vertically inherited genes, diminishing the likelihood that they are due to errors in the metagenome assembly. However, we found 340 HGT “islands”, comprising 797 Bacteria-to-Archaea HGT genes (on average, 31.84% of the Bacteria-to-Archaea HGT genes, SD=11.63). These islands contained up to 8 genes (mean 2.77, SD 1.09). 48 of these islands contained genes belonging to the same COG category, suggesting that they may be operons. We studied these cases in detail. Some of these islands were composed of the same transferred gene duplicated in tandem, such as GCA016840465.1_JPJMNJDP_02194 and GCA016840465.1_JPJMNJDP_02195 from Desulfobacterota to Baldrarchaeum (bcrC) or GCA037305845.1_JPIBEIIO_00281 and GCA037305845.1_JPIBEIIO_00282 (CRYZ) from Patescibateria to Jordarchaeia. An illustrative example of putative operons is GCF008000775.2_JEDDKEKE_02754 and GCF008000775.2_JEDDKEKE_02755, which are excinuclease ABC subunit B and A transferred from Firmicutes to the CR-4 lineage, while protein C is GCF008000775.2_JEDDKEKE_00387, or GCA016840425.1_IAHJMLLN_00393, GCA016840425.1_IAHJMLLN_00394 and GCA016840425.1_IAHJMLLN_00395, transfers to the Heimdall-Loki clade, likely from Nitrospirota that are zinc-transport system znuA/B/C.

Last, we set to assess which HGT proteins may have reached the eukaryotic lineage via the Asgard ancestor of the Last Eukaryotic Common Ancestor (LECA). We assessed all 331 trees flagged as HGT (both Asgard-to-Bacteria and Bacteria-to-Asgard, as we were also interested in assessing whether proteins in eukaryotes may be falsely identified as of bacterial origin) which contained at least one eukaryotic sequence, and 28 contained sequences from at least 3 eukaryotic supergroups suggesting they might have been present in LECA^39^. Upon manual inspection of these candidate phylogenies, we did not find any clear case where a gene of bacterial origin was horizontally transferred to Asgard archaea and passed on through the eukaryotic lineage by vertical inheritance. This has interesting implications for eukaryogenesis, as it implies that the proteins of bacterial origin beyond Alphaproteobacteria that are detected in LECA are unlikely to have come from Bacteria-to-Asgard HGT events in the Asgard lineage prior to eukaryogenesis, strengthening the credibility of the implication of more bacterial partners during eukaryogenesis^40^. Interestingly, however, we do find instances where viruses (mainly Nucleocytoviricota) seem to act as mediators for the transfer of genes from prokaryotes to eukaryotes, supporting and expanding previous observations^39^. These viruses, of Algavirales, Imitervirales, Pimascovirales clades, mainly infect microbial eukaryotes of varied phylogenetic affiliations (amoebozoa, and other protists).

## Discussion

Altogether, our findings depict a high dynamism in terms of content and size of Asgard archaeal genomes and provide the first comprehensive assessment of gene duplication, loss, and HGT within this recently discovered and relevant clade. We reconstructed a large and open Asgard archaeal pangenome, which likely reflects the clade’s immense functional and ecological diversity and suggests a vast, untapped genomic diversity. While acknowledging the current dominance of metagenome-assembled genomes (MAGs) and the scarcity of cultured representatives, which may lead to incorrectly inferred absences due to genome incompleteness, we anticipate the pangenome will likely remain open even with a greater diversity of culture-based genomes.

We identified a core set of 14 Asgard signature proteins (ASPs) widespread in the clade and rarely found in prokaryotic out-groups. Significantly, these ASPs are enriched in endocytosis, a function relevant to eukaryogenesis^41,42^. Notably, several ASPs are annotated with functional domains shared with eukaryotes and were previously considered Eukaryotic Signature Proteins (ESPs)^2^, indicating that Eukaryotes inherited some of the distinctive Asgard archaeal features.

Our comprehensive phylogenomic analysis revealed high gene duplication rates across all lineages—a finding atypical for prokaryotic genomes. Indeed, Asgard archaea exhibited duplication rates intermediate between their closest archaeal relatives (Thermoproteota) and eukaryotic unicellular free-living heterotrophs. This underscores the “stepping-stone” nature of Asgard archaea, even in this genomic trait. Similar to eukaryotes, Asgard archaeal duplications preferentially affected specific gene families, showed lineage-specific patterns, often occurred in tandem, and, alongside gene loss, emerged as a primary driver of genome size. Among the lineages with the highest duplication rates were Heimdallarchaeia (considered the closest sister group to eukaryotes^19^), Lokiarcheia and Sifarchaeia.

We also assessed intra-Asgard transfer, uncovering an intricate network superimposed on a strong vertical inheritance backbone. Intriguingly, genes encoding ESPs were more frequently transferred than other genes, which could explain their previously observed patchy phylogenomic distribution^2^. We estimated that 2-13% of the genes in Asgard genomes were involved in inter-domain HGT with bacteria. This phenomenon is pervasive across the entire Asgard phylogeny and involves bacterial partners that co-occur with Asgard archaea in metagenomic samples. HGT rates are lower to those found in Thermoproteota (when normalizing for genome size), a trend driven by a lower transfer rates with other Archaea (including Thermoproteota), suggesting HGT alone does not account for increased genome sizes in Asgard archaea. The functions of acquired bacterial genes were highly diverse and lineage-specific, predominantly metabolic, and were found to be peripheral in protein-protein and metabolic networks, interspersed with genes of vertical inheritance This pattern suggests a piecemeal integration into established pathways, perhaps replacing ancestral archaeal proteins, rather than the wholesale addition of new functional modules.

In summary, our work provides a detailed survey of genome content dynamics in Asgard archaea, defines the signature proteins of this important clade, and uncovers a rich dynamism of gene family duplication that is atypical for prokaryotes but reminiscent of processes observed in eukaryotic organisms. These insights offer crucial mechanistic details about the genomic evolution of this pivotal lineage and its implications for understanding the origins of eukaryotic complexity.

## Methods

### Data retrieval and annotation

From GTDB r226 we compiled a set of 32 genomes, covering all Asgard family-level lineages *sensu* GTDB and chosen for their best assembly quality parameters (applying the score of CheckM completeness - 5*CheckM contamination), except in the cases in which closed genomes were available, which were prioritized over metagenomes as they served as a baseline that the findings of HGT were not due to contaminating sequences in the metagenomes. We generated a second dataset of Asgard archaea, which was a random set of equal size and same lineage-based representation for quality control of the pangenomic measures. Taxonomies were assigned *sensu* GTDB.

We additionally retrieved representatives of the Thermoproteota superphylum (“p Thermoproteota” *sensu* GTDB), one per order-level lineage, as well as 1 representative per phylum for the rest of the prokaryotic phyla. This generates a dataset of 195 genomes in total.

To avoid biases in genome annotation, we re-annotated all genomes using Prokka v.1.14.6^43^ with the flags --kingdom Archaea (or --kingdom Bacteria, when necessary) and --metagenomes, as Prokka is a reliable and widely used prokaryotic genome annotator. We adapted the headers of the proteome files, changed the NCBI taxID when the one in the GTDB metadata table was too broad or conflicting, and uploaded them to PhylomeDB^30^ via the Phylomizer.

We annotated the Gene Ontology (GO) terms and domains associated with each protein using InterProScan v5.39-77.0^44^ (filtering for score ≤ 1E-3), and the COG categories and KEGG KOs with the COG HMMs^45^ and KoFamScan (https://github.com/takaram/kofam_scan), respectively. To perform functional assessments across the Asgard pangenome, we also annotated the orthogroups by compiling the annotations of their member proteins following the pipeline in^39^. In brief, we followed the strategy followed in eggNOG, which takes into account the proportion of the KO and COG within the set of sequences and its overrepresentation with respect to the background KOs distribution (the whole set of annotated proteins).

When querying for ESPs, we performed a manual curation of InterProScan results. To assign an ESP to a given OG (for example, for the results in Figure 1) we took the OG where the majority of the proteins from that ESP were grouped (or several in case of ties).

We took the annotations from the Asgard proteins for the consensus, to keep the inferred function as close as possible to the actual one performed in Asgard cells, except for when looking for integrase domains, where we took into account Thermoproteota annotations as well.

Last, we taxonomically downsampled the list of Free-Living Unicellular Heterotroph (FLUH) proteomes from Bernabeu et al.^39^, by keeping a proportion of proteomes per supergroup to a total of 30.

The list of genomes, along with associated metadata, can be found in Supplementary Data 1.

### Selected thresholds for Asgard Signature Proteins

We defined signature proteins of the Asgard clade (which we termed Asgard Signature Proteins or ASPs) as those common within Asgard genomes and rare in the outgroups. We determined the abundance thresholds as follows. First, we defined the outgroup abundance threshold at 15% which is the upper limit defined for the ‘cloud’ component of a pangenome as defined in^27^. For the presence within Asgard archaea, we considered the lower limit of the extended core (90%) as too strict, given the high genome incompleteness within the set (see **Results**). Therefore, we analyzed the prevalence of housekeeping protein families that can be thought to be core, in the sense that they perform essential functions for the cell. For this, we annotated the orthogroups by KEGG KOs and COG categories and assessed the prevalence within the Asgard group of protein families annotated as the COG category J (Translation, ribosomal structure and biogenesis) and of the orthogroups annotated as the KOs for the ribosomal proteins in archaea (BRITE for ko03011). We selected a threshold of prevalence of 86%, as this is the median prevalence for the COG category J and similar to the median for ribosomal proteins (87%). This resulted in 14 orthogroups selected as Asgard Signature Proteins.

### Alternative species tree reconstruction

We inferred branch lengths of the Asgard species tree by retrieving the markers for all archaeal species from GTDB, aligning them with MAFFT-linsi v7.407^46^, trimming with trimAl v1.4.rev15^47^, and reconstructing the species tree with IQ-TREE v1.6.9^48^ (--models DCmut,JTTDCMut,LG,WAG,VT), both constraining the topology to the current consensus (see ref.^19^) for the main lineages or to the GTDB topology, resulting in an alternative topology to benchmark results.

### Asgard archaeal pangenome assessment

We performed an orthogroup clustering with OrthoFinder v.2.5.4^49^ using BLAST^50^ as an aligner. To assess pangenome metrics, such as Chao’s lower bound, Heaps estimate, and the binomial mixture models we employed the R package micropan^51^.

We defined Asgard signature proteins (ASPs) as those OG common (present in ≥ 86% of the proteomes) in Asgard archaea but rare (present in ≤15% of the proteomes) in other prokaryotes (see Methods). We mapped ASPs onto the Protein-Protein Interaction Network (PPI), obtained with STRING v12.0^29^, of the seed genome with the highest number of ASPs (*Ca.* P. syntrophicum) using STRING web server.

To perform functional enrichments, we collapsed the annotations at orthogroup level following the aforementioned pipeline.

### Genome dynamics across Asgard archaea

We constructed a metaphylome consisting of a collection of phylomes employing one seed per major lineage. For the phylome analysis, we took the 60 representative genomes of Asgard archaea and Thermoproteota as the dataset and selected 10 seed genomes per phylum that covered the diversity of the phyla. For the set of FLUHs, we selected 4 seed genomes with 30 the aforementioned 30 representatives.

The phylome reconstruction pipeline was applied as defined elsewhere (see ref.^30^ for full details): In brief, for each of the genes contained within each of the seed genomes, we performed a protein BLAST^50^ search against all proteomic data to retrieve proteins with significant similarity (defined as e-value < 1e-05, continuous overlap over 50% of the query sequence). We then took the top 150 hits per protein and generated an MSA with three different methods (MUSCLE V3.8.551^52^, MAFFT v7.407^53^ and KALIGN v2.04^54^), in forward and reverse, and combined the resulting six alignments into one consensus alignment with M-Coffee^55^. This alignment was then trimmed with trimAl v1.4.rev15^47^, with –gappyout. These alignments were then used to reconstruct the phylogenetic tree of each protein family with IQ-Tree v3.0.1^48^, for which we selected the final Maximum Likelihood (ML) tree using the best model selected based on the Bayesian Information Criterion (BIC) and support was calculated with ultrafast bootstrap under 1000 repetitions.

We then ran the species overlap algorithm following the PhylomeDB pipeline^30^. We employed both topologies for the prokaryotic dataset and the species tree as inferred by Duptree^56^ for the eukaryotic set.

For the reconciliation algorithm, which we ran for the prokaryotic set as we infer HGT to play a significant role in its genome dynamics, we built gene trees of the OGs from OrthoFinder that had sequences from more than 4 species following the tree reconstruction pipeline in ref.^39^ (which is an update on ref.^30^) ran AleRax^31^ v1.3.0 using as input the distribution of bootstrap trees (as supplied by IQ-TREE) and the generated species tree as backbone, estimating rates per family.

### Evidence of selection

Within tandem duplications, we looked for signs of positive selection by assessing the dn/ds ratio. For this, we built codon alignments and assessed the dn/ds ratio employing the python library biopython v1.86. Cases where the sequence had “N” or where there were no synonymous mutations were discarded from the analysis.

### Mobile Genetic Elements

We took the InterProScan annotation of the Asgard and Thermoproteota proteomes and assessed the prevalence of signatures related to Mobile Genetic Elements (MGEs). For this we took as reference the list of InterPro domains employed by proMGE^57^. We took the orthogroups for which at least 50% of the proteins with annotations were annotated (score <= 1E-3) with a given domain within this list. We then assessed the number of copies per node of these orthogroups by doing the sum of copies of each orthogroup.

### Inter-domain HGT assessment

We performed a similarity search with BLAST 2.11.0^50^ of the proteomes against a custom-made database (BROADDB_V2), which comprises all species representatives from GTDB r207^5^, viral sequences from RVDB 25.0^58^, and eukaryotes from UniRef v2023_01^59^ and EukProtV1^60^. We chose not to employ NCBI NR as it does not provide a sufficient representation of the Asgard genomes. We employed GTDB taxonomy for prokaryotes as it allows a more comprehensive, robust and fine-grained taxonomic analysis as compared to NCBI taxonomy.

We analyzed the BLAST results with HGTector v2.0b3^61^, which systematically looks for hit distribution patterns incongruent with vertical evolution, given a series of hierarchically defined evolutionary categories. We ran the algorithm setting the “self” group as Asgard archaea and the “close” as Archaea, therefore allowing us to focus on detection of inter-domain HGT events. The output statistics can be found in Supplementary Data 5 and the output of HGTector in the data repository.

For candidate transfers selected by HGTector, we retrieved the best 300 BLAST hits with an e-value cut-off of 0.001 and used them to reconstruct a gene tree following the pipeline in^39^.

When the selected set of homologous sequences was saturated with eukaryotic sequences (that is, if they comprised >=80% of the non-Asgard sequences) or Asgard sequences, we performed a subsampling of these clades by selecting the first hit per major eukaryotic group as defined in EukProt taxonomical framework^60^, then retrieved all hits from those organisms that had a lower e-value than the lowest e-value for a non-Asgard prokaryotic sequence for that query. We kept the non-eukaryotic sequences from the BLAST that passed the threshold, the subsampled eukaryotic sequences, and up to 250 non-eukaryotic sequences passing the threshold performed in a prokaryotic-only BLAST. This allows us to have taxonomically-informative insight into the distribution of the sequences across eukaryotes whilst allowing us to analyze the distribution of more distant prokaryotic homologues.

We further analyzed the resulting gene trees our custom script (HGT_detection.py) (see below for the pseudo-code) and with Abaccus v1.2^62^, which identifies taxonomical “jumps” within gene trees that do not follow the species tree and therefore further helps discern putative HGT events. For this, we used the taxonomy *sensu* GTDB for prokaryotes.

There are several methods to root a phylogenetic tree in absence of a clear outgroup. We tested the number of HGT events called by rooting the trees by midpoint and Minimal Ancestral Deviation (MAD), which resulted in a 94% overlap in the genes detected as HGT (2,946), 2% MAD-only (54) and 5% midpoint-only (244). Given the small difference, we decided to proceed with MAD rooting as implemented in toytree^63^.

Additionally, we retrieved the support of the branch of the transfer (e.g. the branch subtending the acceptor Asgard clade and the putative non-archaeal donor clade). We only kept cases when this support was ≥50%, or, if the node was poorly supported, cases where the sister of the transfer branch contained ≤ 10% of archaeal sequences were also considered, even if the specific donor clade could not be ascertained. To ensure that branch support was not affecting the conclusions of the analysis, we built a subset of well-supported (≥ 70% support) HGT cases in which all analyses were also run.

Last, we manually curated the HGT results by filtering out incongruencies and false positives resulting from trees saturated with eukaryotic sequences. We performed this step for all proteins flagged by HGTector, whether they were picked up by Abaccus or not.

To assess the directionality of the transfer (Asgard-to-bacteria or bacteria-to-Asgard), we assessed whether there were Asgard sequences in sister2, if there were, Asgard-to-bacteria was assumed.

### HGT detection pipeline

To do so, we first numbered the internal nodes in the Asgard tree . Then, we parsed the gene trees with an in-house script (HGT_detection.py) based on ete3^64^, using the following criteria:

1. Trees were rooted with Minimum Ancestor Deviation (MAD)^65^.
2. The start node was established as the one subtending the largest Asgard monophyletic clade containing the seed protein. In assessing monophyly, we considered eukaryotes as a *de facto* Asgard lineage stemming from Heimdallarchaea. The most recent common ancestor (MRCA) of the Asgard lineages present in the clade is assumed as the acceptor Asgard lineage.
3. Then, the sister clade of the start node was analyzed:

a. If the fraction of non-Asgard Archaeal sequences in the sister clade was ≥ 10%, the tree was labeled as a False Positive. This means that either vertical inheritance cannot be ruled out, or that an Archaeal-to-Asgard HGT cannot be ruled out, which is also beyond the scope of this analysis since we are focusing on inter-domain HGT.
b. The donor was set as the MRCA of the non-Asgard sequences of the sister clade. In cases where the MRCA is a broad category (e.g “Bacteria”), we established the predominant phyla as the putative donor lineage. For cases with low support (<= 50) we marked the donor as “uncertain”.
4. Additionally, taxonomy information of the sister clade to the (start+sister) branch (sister2) was retrieved.
5. We assessed the percentage of non-Asgard sequences in the tree that were of eukaryotic origin. In cases where this percentage ≥ 80% were labeled eukaryote-saturated and submitted to the subsampling mentioned above and the subsampled gene tree was re-analyzed.

After this analysis, we additionally flagged as FP events for which support was low (≤ 50) and sister2 contained ≥ 10% non-Asgard archaeal sequences. These criteria are very restrictive and likely result in an underestimation of the true number of HGT events: namely, we will miss inter-domain HGT and the 10% of Archaeal sequences in the sister needed to label a tree as a false positive will lead to true HGTs being missed due to secondary transfers of the gene family from bacteria to other archaea. However, this ensures that the trees that are picked up as HGT are true HGT events.

One of the limitations of the analysis of acceptor nodes of horizontally transferred genes is the use of Maximum Parsimony. There are methods better suited for this (e.g reconciliation, as used for the estimation of duplication, transfer and loss rates), but these become computationally prohibitive given that running time scales cubic to the number of species, and given that the search is done against a broad database of almost 500.000 genomes. Moreover, for the purpose of analyzing the tempo of transfers across Asgard phylogeny (that is, whether there are ‘hotspots’ or it is more widespread) we believe a more simplistic assumption like Maximum Parsimony is sufficient for this purpose.

### Tandem duplications and HGT islands

To infer tandem duplications we used a window-based approach to query the GFF files where we looked for 2 or more genes belonging to the same OG being either directly adjacent or interspaced by another gene, which we applied with a custom R script (Tandem_Duplications.R). For HGT islands, we also queried the GFFs to find HGT genes gene immediately adjacent or interspaced by a vertically inherited gene to reconstruct “islands”. We visualized it with the R package gggenes 0.5.1 (https://github.com/wilkox/gggenes/tree/master).

### Functional enrichment analysis

We performed GO term enrichment analyses using the Bioconductor package topGO (https://git.bioconductor.org/packages/topGO), with the method “weight01” and using the Fisher’s exact test, correcting for multiple testing and keeping results with a False Discovery Rate (FDR ≤ 0.15). For KEGG pathway enrichment, we employed the package ClusterProfiler^66^ with the same statistical criteria except in the tandem duplications, where we cut to FDR ≤ 0.1 for visualization purposes. We visualized the results with the R package ggpubr (https://github.com/kassambara/ggpubr/) and ggtree/ggtreeExtra^67,68^.

### Protein-protein interaction networks

We further analyzed the functional relationship between transferred genes by assessing the network of protein-protein interactions (PPI). Since none of the analyzed Asgard genomes are available in the database, we first submitted them to be added to STRING^29^ v.12.0 by making use of the tool developed in the latest version^69^. We then uploaded the sequences of the transferred genes to the web server and ran the analysis using the newly uploaded proteome as target organism. We downloaded the results, performed analysis with the R implementation of iGraph (https://github.com/igraph/rigraph) and visualized the networks with the R package ggnetwork (https://journal.r-project.org/archive/2017/RJ-2017-023/index.html).

## Data availability

BioProject IDs for the genomes are available in Supplementary Data 1. For the representative set of Asgard genomes, the publications (if any) associated with the assemblies are as follows: GCA_001940655.1^2^; GCA_005191415.1 and GCA_005191425.1^70^; GCA_014729935.1^71^; GCA_015523565.1^72^; GCA_024720975.1^73^; GCA_021513695.1^18^; GCA_001940665.2^74^; GCA_027061945.1, GCA_027068385.1, and GCA_026993975.1^75^, GCA_016292335.1^76^; GCA_029856635.1^19^; GCA_040225955.1 and GCA_029838215.1^77^, GCA_030638685.1 and GCA_030671945.1^78^; GCA_038858525.1^79^; GCA_037305845.1 and GC_037308085.1^80^; GCF_008000775.2^7,81^ All input proteomes and output files for all steps can be found in Zenodo (10.5281/zenodo.15789115)

## Code availability

All described scripts are available on Github (https://github.com/Gabaldonlab/Asgard)^82^.

## Acknowledgements

We want to thank Moises Bernabeu for the BroadDB database and Giacomo Mutti for help with using AleRax. TG group acknowledges support from the Spanish Ministry of Science and Innovation (grant numbers PID2021-126067NB-I00 and PLEC2023-010225), cofounded by ERDF “A way of making Europe”, as well as support from the Gordon and Betty Moore Foundation (grant number GBMF9742); “La Caixa” foundation (grant number LCF/PR/HR21/00737 and CI23-20260), Fundació La Marató de TV3 (202328-31), AECC (PRYGN234923GABA and 290059), and Instituto de Salud Carlos III (CIBERINFEC CB21/13/00061-ISCIII-SGEFI/ERDF and Proyectos de Desarrollo Tecnológico en Salud (DTS) (grant number DTS25/00141).

## Author contributions

SM-M performed all bioinformatics analyses, wrote the necessary code, and produced the visualisations. TG conceptualised the project, obtained funding and supervised the project. All authors contributed to the design of the analytical steps, to the interpretation of the results, and to writing the manuscript.

## Competing Interests

The authors declare no conflicts of interest.

## Notes

### Competing Interest Statement

The authors have declared no competing interest.

### Summary of Updates

Some analysis were updated based on revision by peers

