## Supplementary text for "Phylogenomics of Asgard archaea reveals a unique blend of prokaryotic-like horizontal transfer and eukaryotic-like gene duplication"

08034 Barcelona, Spain.

2) Institute for Research in Biomedicine (IRB Barcelona), The Barcelona Institute of Science and Technology, Baldiri Reixac, 10, 08028 Barcelona, Spain

3) Catalan Institution for Research and Advanced Studies (ICREA), Barcelona, Spain

4) CIBER de Enfermedades Infecciosas, Instituto de Salud Carlos III, Madrid, Spain

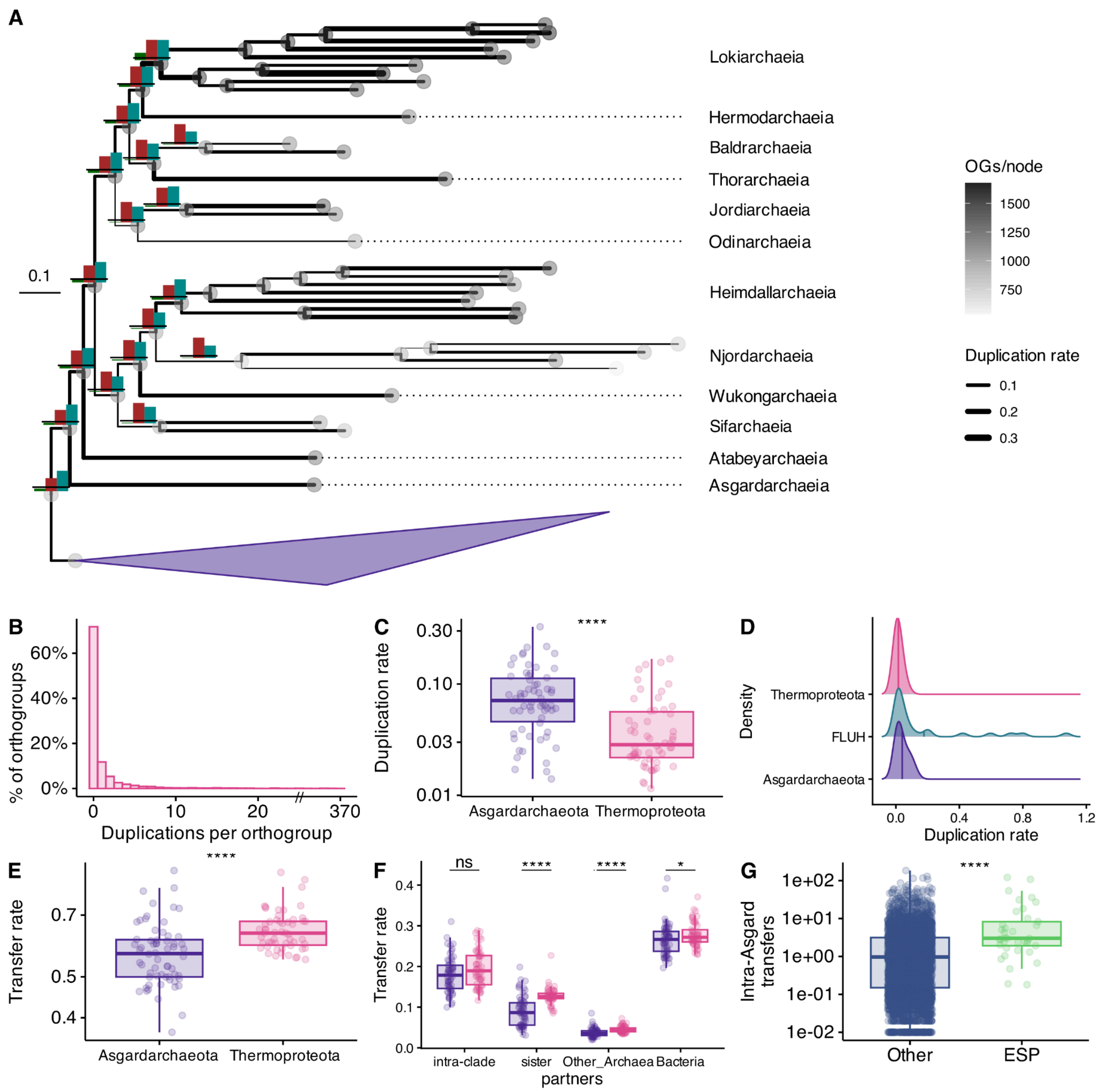


**Supplementary Figure 1: Gene gain-loss dynamics across Asgard archaea for the alternative topology.** (A) Gain and loss dynamics in Asgard archaea. The total number of OGs per proteome is displayed by circle color in each node. Duplication rate (number of duplication events relative to the inferred number of families present in the node) is represented by the branch width. Number of duplications (green), transfers (blue) and losses (red) displayed at internal nodes in absolute number. Horizontal bar indicates 100 events as scale. (B) Histogram of sum of duplication events per orthogroup across Asgard nodes. Axis cut at 30, we artificially display the tick as the highest value rounded to the multiple of 10.(C) Duplication rates per node of Asgard (n=62) and Thermoproteota (n=54) gene families, statistical significance assessed with two-tailed Wilcoxon rank-sum test, p-value 1.85e-06. (D) Duplication rates, inferred by species overlap, of Asgard, Thermoproteota and FLUHs. (E) Transfer rates per node of Asgard (n=62 nodes) and Thermoproteota (n=54 nodes) gene families, statistical significance assessed by two-tailed Wilcoxon rank-sum test, p-value 4.9E-6 (F) Transfer rates in Thermoproteota (n=54) and Asgard archaea (n=62) nodes, separated by donor, statistical significance assessed by pairwise two-tailed Wilcoxon rank-sum tests, p-values 0.0665 (intra-clade), 9.03E-09 (sister), 2.51E-06 (other archaea), 0.0425 (bacteria) (G) Intra-Asgard transfer events on ESPs compared to the rest of gene families (n=37, 4308 gene families, respectively). Permutation test, p-value 2.2E-16. In the plots with significance values, asterisks * indicate p-values, * (cut-off 0.05) ** (0.01), *** (0.001) **** (0). For the box plots in 2C, E-G, in each boxplot, the central line indicates the median, the lower and upper limits of the rectangle indicate the first and third quartiles, respectively and the whiskers extend to the minimum and maximum value of 1.5 times the interquartile range.

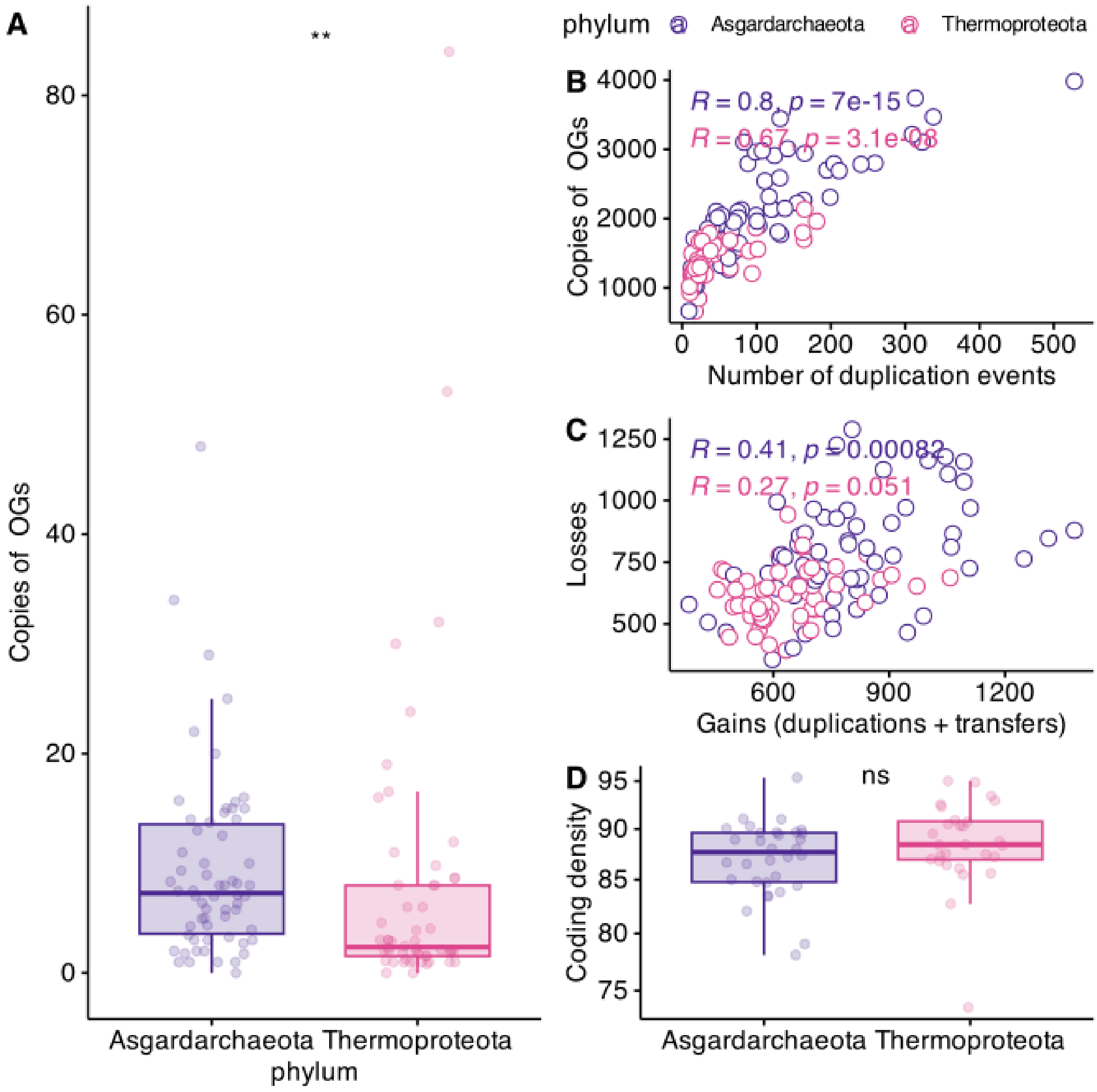


**Supplementary Figure 2:** (A) Number of copies within orthogroups associated with integrase domains in Asgard (n=62) and Thermoproteota (n=54) nodes. Statistical significance obtained by two-tailed Wilcoxon rank-sum test, p-value 1.4628E-3. (B) Correlation between total copies of proteins for all orthogroups and number of duplication events per nod in Asgard and Thermoproteota nodes. (C) Correlation between loss events and gain events (duplications + transfers) in Asgard and Thermorproteota nodes. (D) Comparison between coding density in Asgard (n=32) and Thermoproteota (n=28) genomes. Statistical significance obtained by two-tailed Wilcoxon rank sum test, p-value 9.87E-2. In each boxplot, the central line indicates the median, the lower and upper limits of the rectangle indicate the first and third quartiles, respectively and the whiskers extend to the minimum and maximum value of 1.5 times the interquartile range. . In the plots with significance values, asterisks * indicate p-values, * (cut-off 0.05) ** (0.01), *** (0.001) **** (0).

### Eme et al. (2023) topology

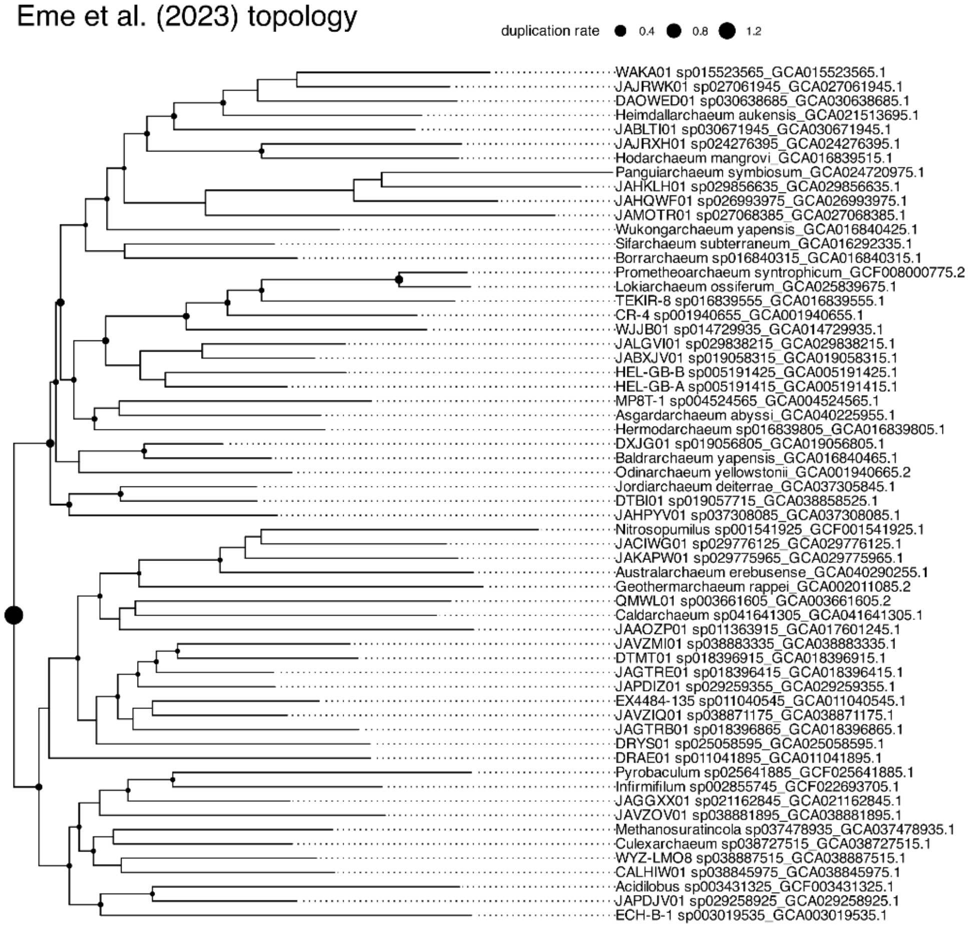


### GTDB topology

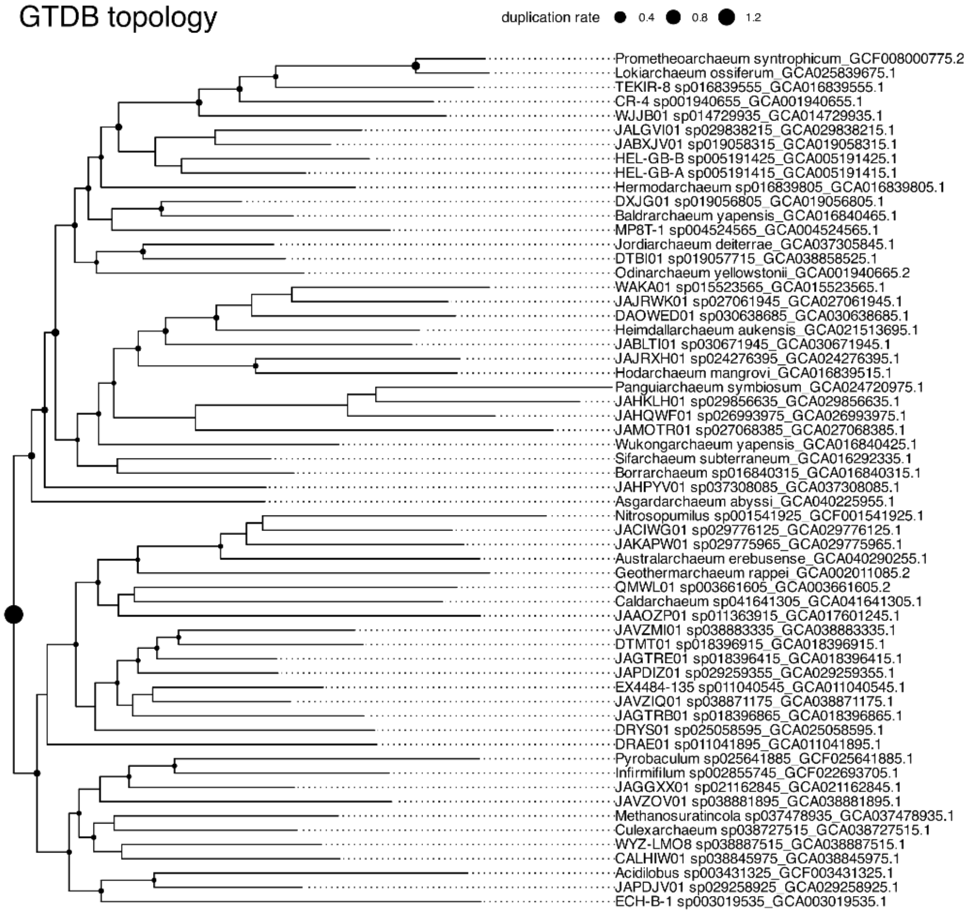

**Supplementary Figure 3: Duplication rates inferred with the species overlap algorithm.** Backbone tree of Asgard and Thermoproteota. Duplications shown for internal nodes that appear in the phylome data. Top: Eme et al (2023) topology. Bottom: GTDB topology.

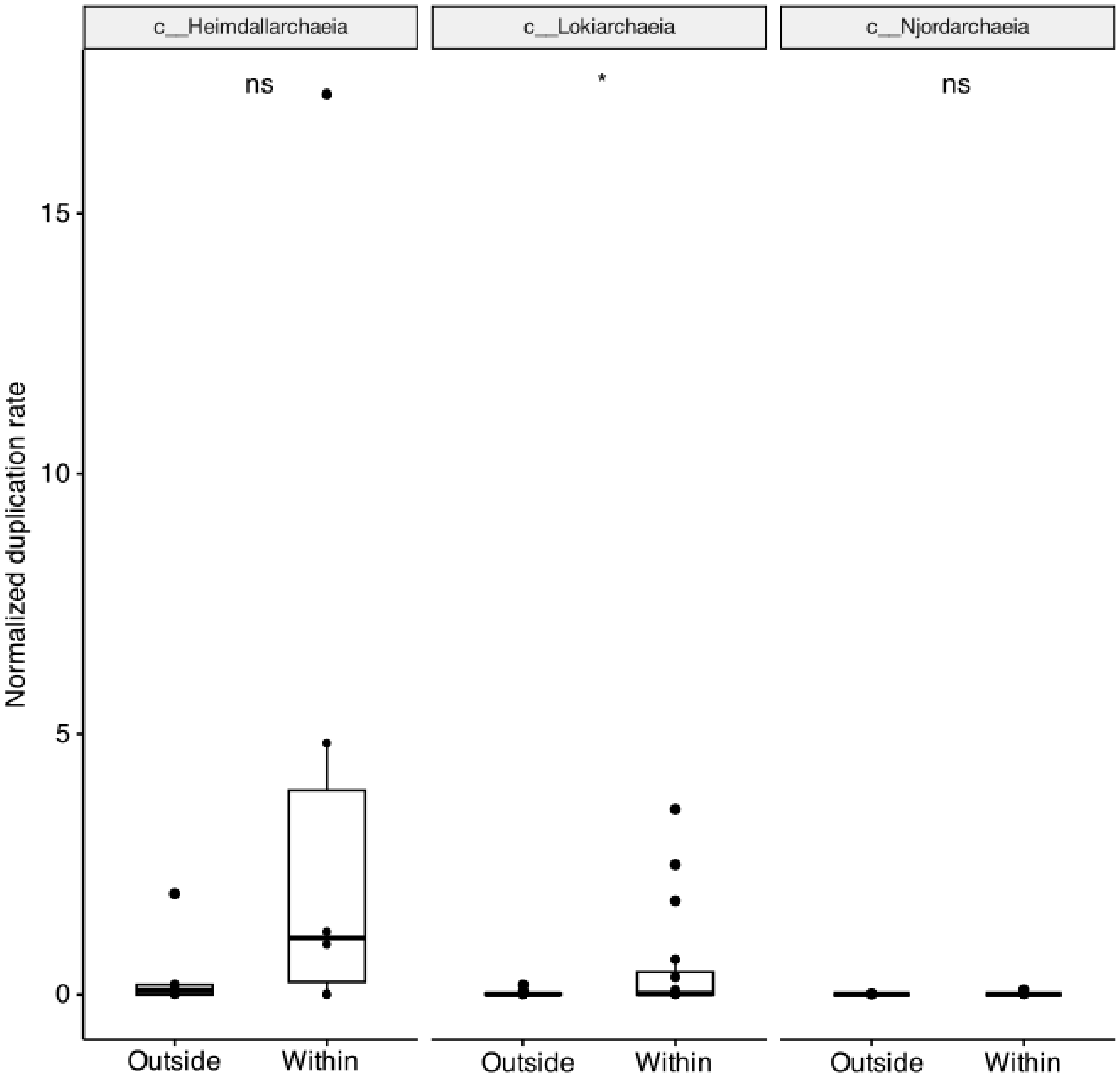


**Supplementary Figure 4: Comparison of duplication rates within- and outside-lineage for signature gene families.** Signature gene families are defined as those common within the lineage but rare outside it. Statistical significance as defined by paired two-tailed Wilcoxon rank-sum tests, p-values 0.2 (Heimdallarchaeia, n=6 orthogroups), 0.011 (Lokiarchaeia, n=20 orthogroups), 1 (Njordarchaeia, n=10 orthogroups). In each boxplot, the central line indicates the median, the lower and upper limits of the rectangle indicate the first and third quartiles, respectively and the whiskers extend to the minimum and maximum value of 1.5 times the interquartile range. Asterisks * indicate p-values, * (cut-off 0.05) ** (0.01), *** (0.001) **** (0).

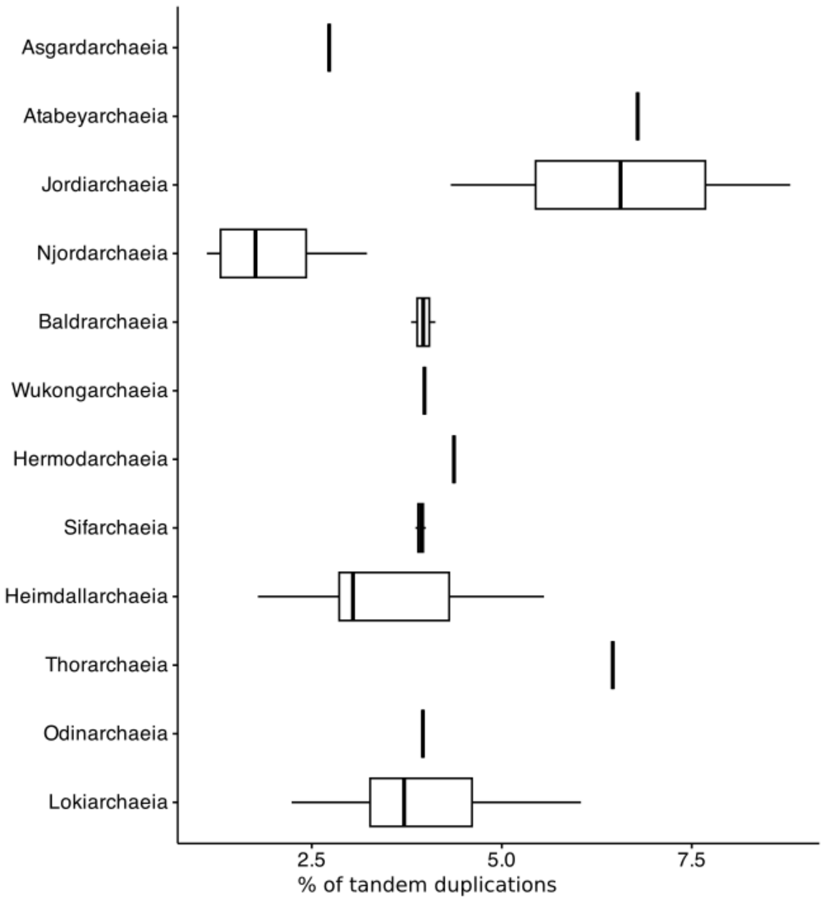

Asgardarchaeia
Atabeyarchaeia
Jordiarchaeia
Njordarchaeia
Baldrarchaeia
Wukongarchaeia
Hermodarchaeia
Sifarchaeia
Heimdallarchaeia
Thorarchaeia
Odinarchaeia
Lokiarchaeia
2.5
5.0
7.5
% of tandem duplications


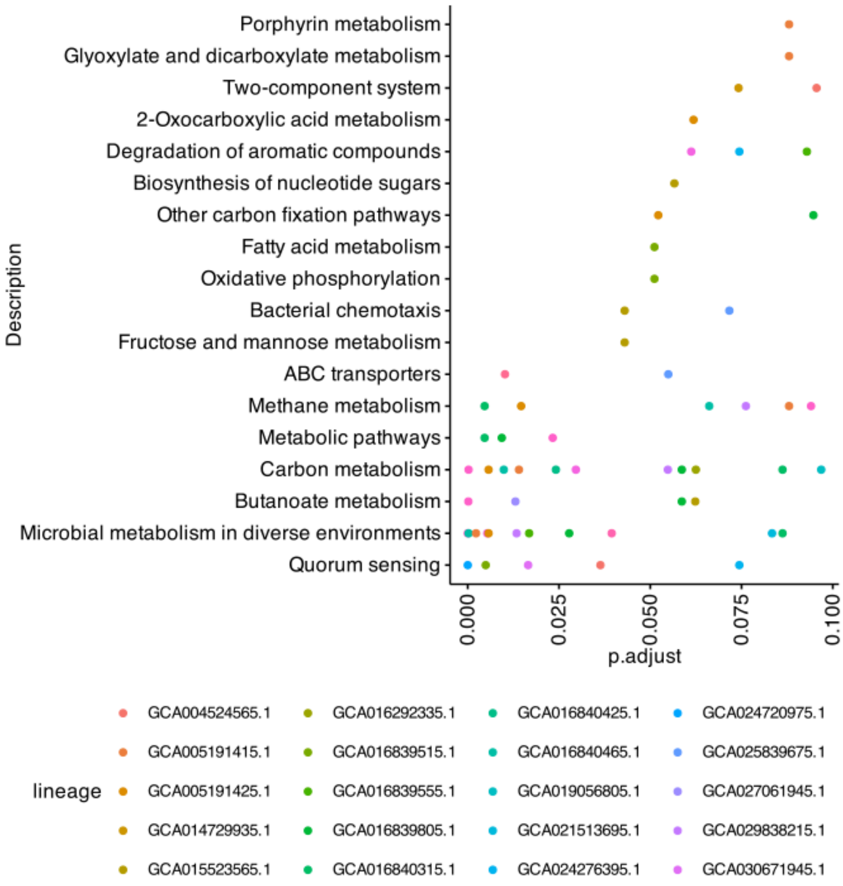

Porphyrin metabolism
Glyoxylate and dicarboxylate metabolism
Two-component system
2-Oxocarboxylic acid metabolism
Degradation of aromatic compounds
Biosynthesis of nucleotide sugars
Other carbon fixation pathways
Fatty acid metabolism
Oxidative phosphorylation
Bacterial chemotaxis
Fructose and mannose metabolism
ABC transporters
Methane metabolism
Metabolic pathways
Carbon metabolism
Butanoate metabolism
Microbial metabolism in diverse environments
Quorum sensing
Description
0.000
0.025
0.050
0.075
0.100
p.adjust
lineage
GCA004524565.1
GCA005191415.1
GCA005191425.1
GCA014729935.1
GCA015523565.1
GCA016292335.1
GCA016839515.1
GCA016839555.1
GCA016839805.1
GCA016840315.1
GCA016840425.1
GCA016840465.1
GCA019056805.1
GCA021513695.1
GCA024276395.1
GCA024720975.1
GCA025839675.1
GCA027061945.1
GCA029838215.1
GCA030671945.1

**Supplementary Figure 5: Tandem duplications in Asgard archaea** (A) Percentage of genes within tandem duplications with respect to the total number of duplicated genes per lineage.In each boxplot, the central line indicates the median, the lower and upper limits of the rectangle indicate the first and third quartiles, respectively and the whiskers extend to the minimum and maximum value of 1.5 times the interquartile range. (B) KEGG pathway enrichment per genome

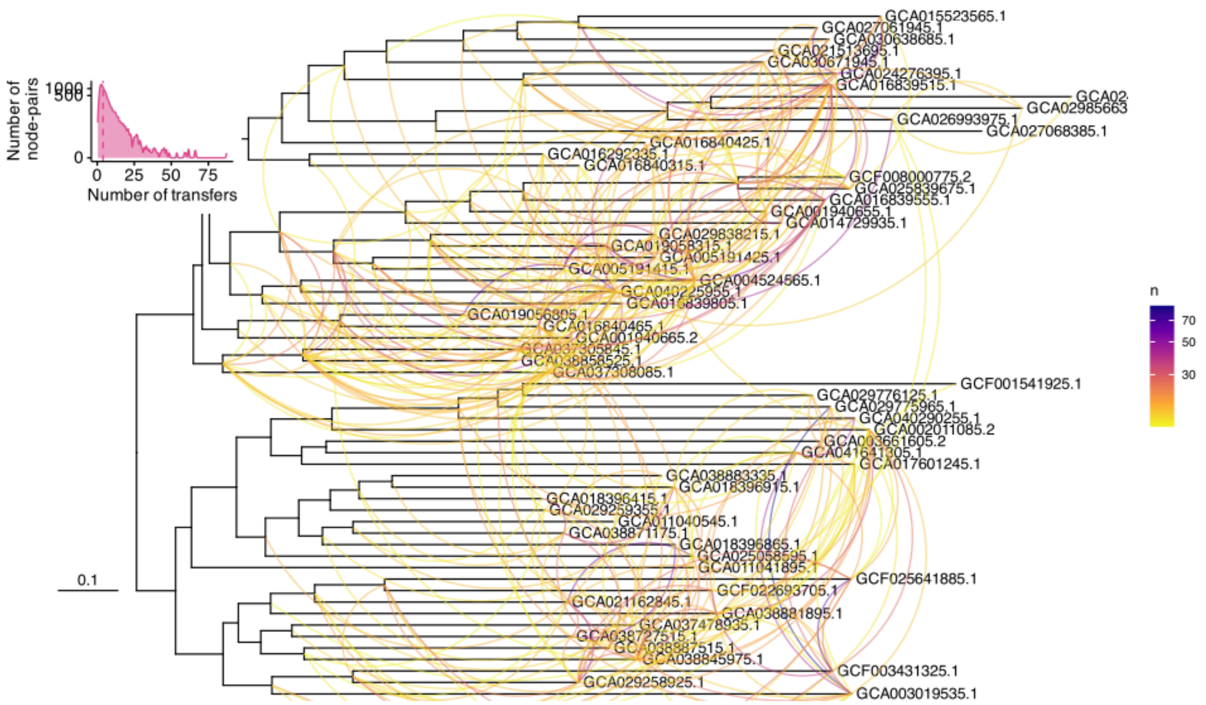


**Supplementary Figure 6:** Network of transfer pairs as inferred by ALERax across Asgard-Thermoproteota. The scale in the tree depicts the number of substitutions per site. Only node-pairs with a number of transfer events higher than percentile 95 are displayed, distribution of the number of transfers per transfer-pairs depicted in the inset plot on the top right.

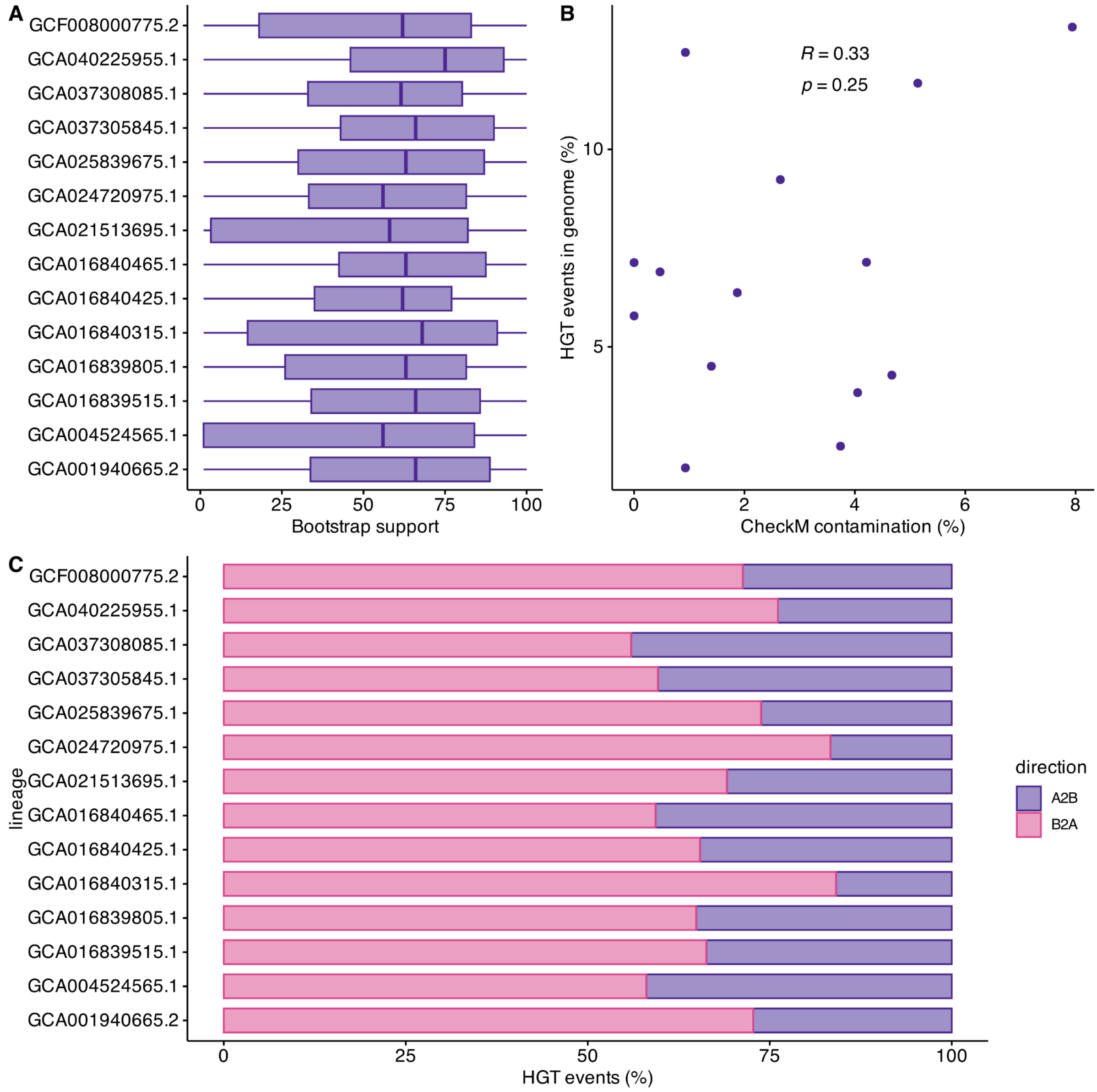


**Supplementary Figure 7:** (A) Distribution of bootstrap support for the node that joins the Asgard acceptor clade with its inferred sister. (B) Correlation between estimated percentage of HGT genes and estimated CheckM contamination per proteome. (C) Directionality of the transfers. Transfers are considered "Asgard to Bacteria" or A2B if the second sister is majoritarily of Asgardarchaeotal taxonomic affiliation, and "Bacteria to Asgard" or B2A if the second sister is majoritarily bacterial. In each boxplot, the central line indicates the median, the lower and upper limits of the rectangle indicate the first and third quartiles, respectively and the whiskers extend to the minimum and maximum value of 1.5 times the interquartile range.

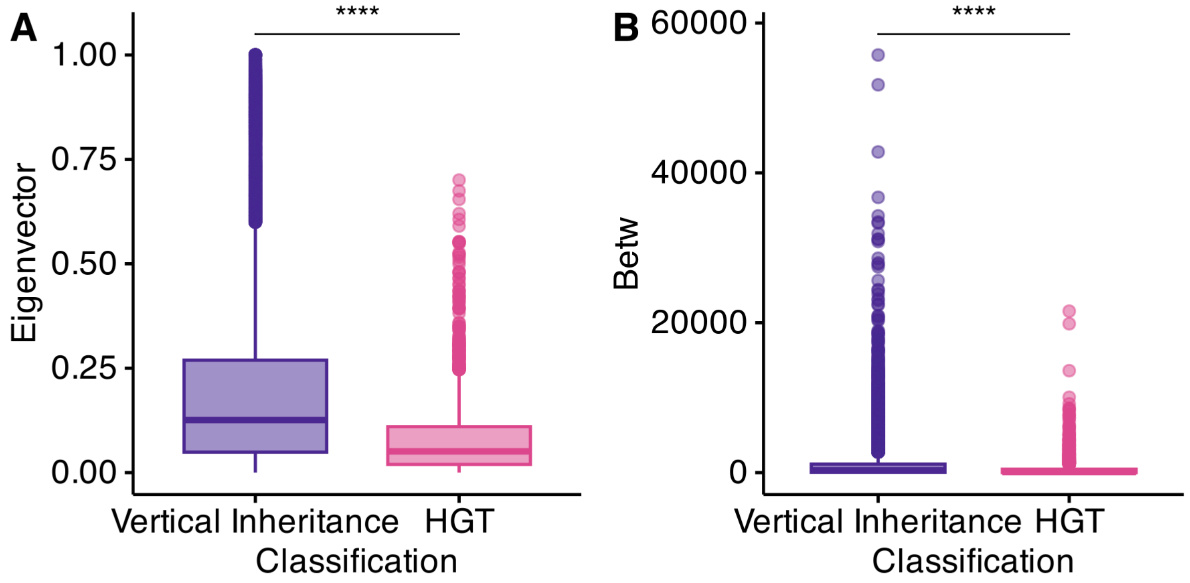


**Supplementary Figure 8: Analysis of alternative centrality measures for HGT proteins (n=2013) and vertically inherited proteins (n=23443,) in Protein-Protein interaction networks.** Values for eigenvector centrality (A) and betweenness (B). In each boxplot, the central line indicates the median, the lower and upper limits of the rectangle indicate the first and third quartiles, respectively and the whiskers extend to the minimum and maximum value of 1.5 times the interquartile range. Significance by two-tailed Wilcoxon rank-sum test, p-values <2.2E-16 for both measures.
